# Hepatocyte nuclear factor-1β recruits the nucleosome remodeling and deacetylase complex to activate gene transcription

**DOI:** 10.64898/2026.10.23.759945

**Authors:** Siu Chiu Chan, Micah D Gearhart, Ziteng Diao, LeeAnn Higgins, Todd Markowski, Annie Shao, Peter Igarashi

## Abstract

Mutations affecting the transcription factor hepatocyte nuclear factor-1β (HNF-1β) are the most common monogenic cause of congenital anomalies of the kidney and urinary tract. The mechanism whereby HNF-1β regulates gene transcription remains unclear. Here, we performed an unbiased proteomic screen and found that endogenous HNF-1β interacts with core subunits of the nucleosome remodeling and deacetylase complex (NuRD) in renal epithelial cells. Genome-wide ChIP-seq studies demonstrated colocalization of endogenous HNF-1β and the core NuRD subunits GATAD2A, CHD4, and MBD3 at 2,776 primarily intergenic and intronic sites in native chromatin from wild-type renal epithelial cells. NuRD co-occupancy at 628 of the sites (22.6%) was reduced in the absence of HNF-1β. Of the 628 HNF-1β-dependent NuRD-occupied sites, 113 sites (18.2%) showed higher ATAC-seq peaks, indicative of open chromatin, and enrichment of histone H3 lysine-27 (H3K27) acetylation, indicative of active enhancers, in wild-type cells compared to HNF-1β-deficient cells. Correlation with transcriptomic profiles in wild-type and HNF-1β-deficient cells revealed that most of the sites were located near genes that are activated by HNF-1β. Knockdown of CHD4 with shRNA resulted in decreased expression of 67 HNF-1β-regulated genes that contained nearby NuRD and HNF-1β co-occupied sites. Collectively, these results demonstrate that HNF-1β recruits the NuRD complex resulting in H3K27 acetylation and chromatin opening and leading to gene activation. Our findings reveal a new mechanism of HNF-1β-dependent gene expression and expand the role of the NuRD complex in gene activation at distal enhancers.

## Introduction

Hepatocyte nuclear factor-1β (HNF-1β) is a homeodomain-containing transcription factor that regulates gene expression in epithelial organs such as kidney, pancreas, liver, and urogenital tract (1). Mammalian HNF-1β consists of 557 amino acids comprising an N-terminal dimerization domain, a central DNA-binding domain, and a C-terminal transactivation domain (2). HNF-1β binds DNA as an obligatory homodimer or heterodimer with the paralogous protein, hepatocyte nuclear factor-1α (HNF-1α) (3). Dimerization is mediated by the N-terminal domain and is dependent on the binding of a dimerization cofactor, pterin-4 alpha-carbinolamine dehydratase 1 (PCBD1) (4). HNF-1β recognizes the consensus DNA sequence 5’-GTTAATNATTAAC-3’ through a bipartite motif comprising a POU homeodomain (POUH) and POU-specific domain (POUS) (5, 6). The C-terminal transactivation domain of HNF-1β mediates the interaction with transcriptional coactivators, such as CBP and PCAF, and corepressors, such as HDAC1 and HDAC2 (7).

HNF-1β plays essential roles in tissue-specific gene expression and the maintenance of tubular epithelial integrity in the organs in which it is expressed (8). In the kidney, the expression of HNF-1β is restricted to tubular epithelial cells in nephrons and collecting ducts. During embryonic development, HNF-1β is expressed in the ureteric bud and comma- and S-shaped bodies that will give rise to collecting ducts and nephrons. In the adult, expression of HNF-1β is maintained in all nephron segments and collecting ducts, where it regulates genes involved in tubular differentiation, solute transport, and cellular metabolism. Human heterozygous mutations of *HNF1B*, encoding HNF-1β, represent the most common monogenic cause of congenital anomalies of the kidney and urinary tract (9). Additional syndromes caused by mutations of *HNF1B* include renal cysts and diabetes, medullary cystic kidney disease, and autosomal dominant tubulointerstitial kidney disease, collectively termed *HNF1B* nephropathy (10). A prominent feature of *HNF1B* nephropathy is the formation of kidney cysts that may phenocopy autosomal dominant polycystic kidney disease (11). Our previous studies have shown that HNF- 1β plays a central role in a transcriptional network that regulates the expression of cystic disease genes, such as *PKD2* and *PKHD1* (*12*). Additional manifestations of *HNF1B* nephropathy include hypokalemia, hyperuricemia, polyuria, and hypomagnesemia (10, 13). Extrarenal features include pancreatic abnormalities, abnormal liver function, and neurodevelopmental abnormalities (9, 10). Genome-wide association studies have linked *HNF1B* to cancers of the prostate, kidney, and ovary (14) .

Despite its clinical importance, the molecular mechanisms underlying HNF-1β-dependent transcriptional regulation remain incompletely understood. Here, we employed rapid immunoprecipitation mass spectrometry of endogenous proteins (RIME) and identified novel HNF-1β-associated proteins in native chromatin. These studies revealed that HNF-1β interacts with multiple core subunits of the nucleosome remodeling and deacetylase complex (NuRD). Multiomic analysis revealed that HNF-1β and NuRD are co-localized on enhancers near HNF-1β target genes in association with increased chromatin accessibility, histone acetylation, and transcriptional activation. Knockdown of the NuRD subunit CHD4 reduced the expression of a subset of HNF-1β target genes. These findings reveal an unexpected role for NuRD in HNF-1β- dependent gene activation at distal enhancers.

## Results

### Identification of HNF-1β interacting proteins by rapid immunoprecipitation mass spectrometry of endogenous proteins (RIME)

To obtain insights into the mechanism whereby HNF-1β regulates gene transcription, we performed liquid chromatography-tandem mass spectrometry (LC-MS/MS) to identify novel proteins that interact with endogenous HNF-1β in native chromatin. We employed a Rapid Immunoprecipitation Mass Spectrometry of Endogenous Proteins (RIME) method to identify endogenous protein complexes by affinity purification at low protein abundance (15). Mouse mIMCD3 epithelial cells, which originated from renal collecting ducts, were cross-linked, nuclei were extracted and digested with micrococcal nuclease, and nuclear lysates were immunoprecipitated with a polyclonal antibody against HNF-1β (Fig. S1A). Coprecipitated proteins were digested on-bead, and the peptides were subjected to LC-MS/MS. Unique peptides were identified using Proteome Discoverer (16). To control for non-specific binding, parallel experiments were performed using nuclear lysates from HNF-1β-deficient mIMCD3 cells in which *Hnf1b* was deleted by CRISPR/Cas9 gene editing (17). Only proteins that were present in two biological replicates in wild-type (WT) mIMCD3 cells and absent in HNF-1β-deficient (KO) cells were selected for further study.

Using RIME and Proteome Discoverer analysis, we identified 484 nuclear proteins that interacted with HNF-1β in WT mIMCD3 cells (Table S1). As validation of the method, we identified HNF-1β itself and known proteins, PCBD1, KPNA2 (Karyopherin subunit alpha-2, Importin subunit alpha-1), and Zyxin, which have previously been shown to interact with HNF-1β (18–20). HNF-1α was not identified since its expression is restricted to proximal tubules, whereas mIMCD3 cells are derived from collecting ducts, further confirming the specificity of the method. Fig. 1A shows representative MS/MS spectra of doubly charged peptides from HNF-1β and the known interacting proteins. The peptides correspond to amino acid residues 94-107, 316-330, 89-104, and 36-54 of HNF-1β, KPNA2, PCBD1, and Zyxin, respectively. The coverage of peptides derived from HNF-1β, PCBD1, Zyxin, and KPNA2 that were identified by RIME in WT mIMCD3 cells ranged from 3.4% to 27.9% (Fig. 1B). In addition to known interacting proteins, we also identified proteins that have not previously been shown to interact with HNF-1β. The novel HNF-1β interactome consists of chromatin remodeling complexes, histone readers, writers, erasers, transcription factors and elongation factors that play important roles in chromatin remodeling and transcriptional regulation (Fig.1C).

**Figure 1.**
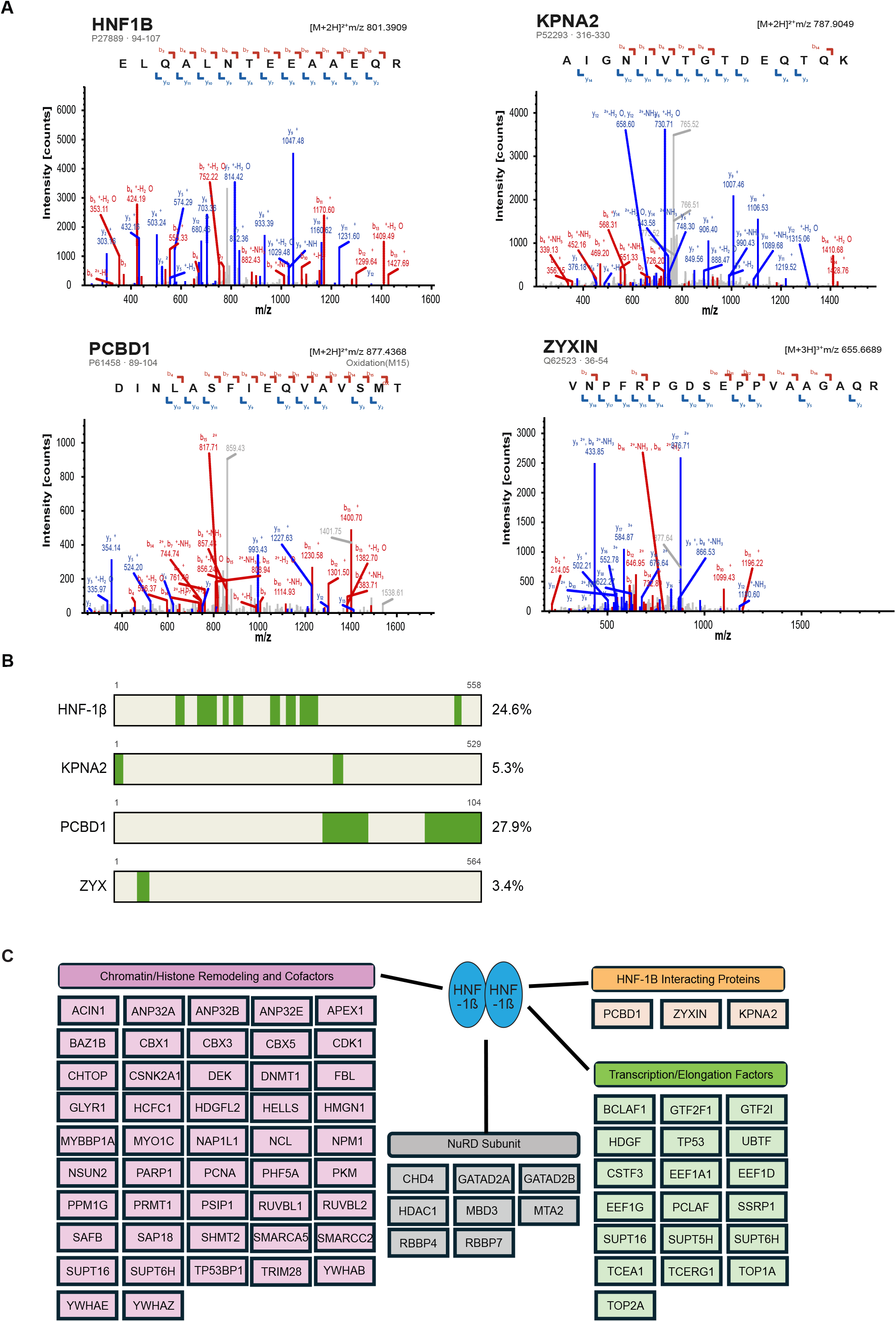
Identification of HNF-1β-interacting proteins by rapid immunoprecipitation mass spectrometry of endogenous proteins (RIME). (A) HNF-1β-interacting proteins were identified by RIME performed on nuclear lysates from wild-type mIMCD3 cells. HNF-1β-deficient mIMCD3 cells were used as negative controls. Representative MS/MS spectra of doubly charged peptides corresponding to HNF-1β and known HNF-1β-interacting proteins, KPNA2, PCBD1, and Zyxin, are shown. Red represents b-fragment ions generated from the C-terminal sequence; blue represents y-fragment ions generated from the N-terminal sequence. Intensity (counts) is plotted against m/z. (B) Coverage of peptides (green) derived from HNF-1β, PCBD1, ZYXIN, and KPNA2 that were identified by RIME in wild-type mIMCD3 cells. (C) Novel HNF-1β-interacting proteins identified by RIME were classified as chromatin remodelers and cofactors (pink), NuRD core components (gray), and transcription/elongation cofactors (green). Results of two biological replicates are shown.

### HNF-1β interacts with core subunits of the nucleosome remodeling and deacetylase complex (NuRD)

Among the novel HNF-1β-interacting proteins, we identified eight distinct core subunits of the nucleosome remodeling and deacetylase complex (NuRD). NuRD subunits CHD4, GATAD2A, GATAD2B, MBD3, HDAC1, MTA2, RBBP4, and RBBP7 were co-immunoprecipitated with an antibody against HNF-1β and identified by LC-MS/MS in WT mIMCD3 cells but not in KO cells. Representative MS/MS spectra are shown in Fig. S1B, and the corresponding peptide sequences and peptide coverage are shown in Fig. 2A-B. A query of the STRING (Search Tool for the Retrieval of Interacting Genes/Proteins) database using a minimum interaction score of 0.7 (high confidence) produced the association network shown in Fig. S2 (21). To validate the interaction between HNF-1β and NuRD subunits, we performed co-immunoprecipitation of endogenous proteins in nuclear lysates from mIMCD3 cells. Western blotting confirmed that HNF-1β was immunoprecipitated in lysates from WT mIMCD3 cells but not in lysates from KO cells (Fig. 2C, left panel). Immunoprecipitation using antibodies against GATAD2A, MBD3, and RBBP4 resulted in co-precipitation of HNF-1β in nuclear lysates from WT cells but not in lysates from KO cells (Fig. 2C, right panel). These findings confirm that HNF-1β interacts with the NuRD complex in the nucleus.

**Figure 2.**
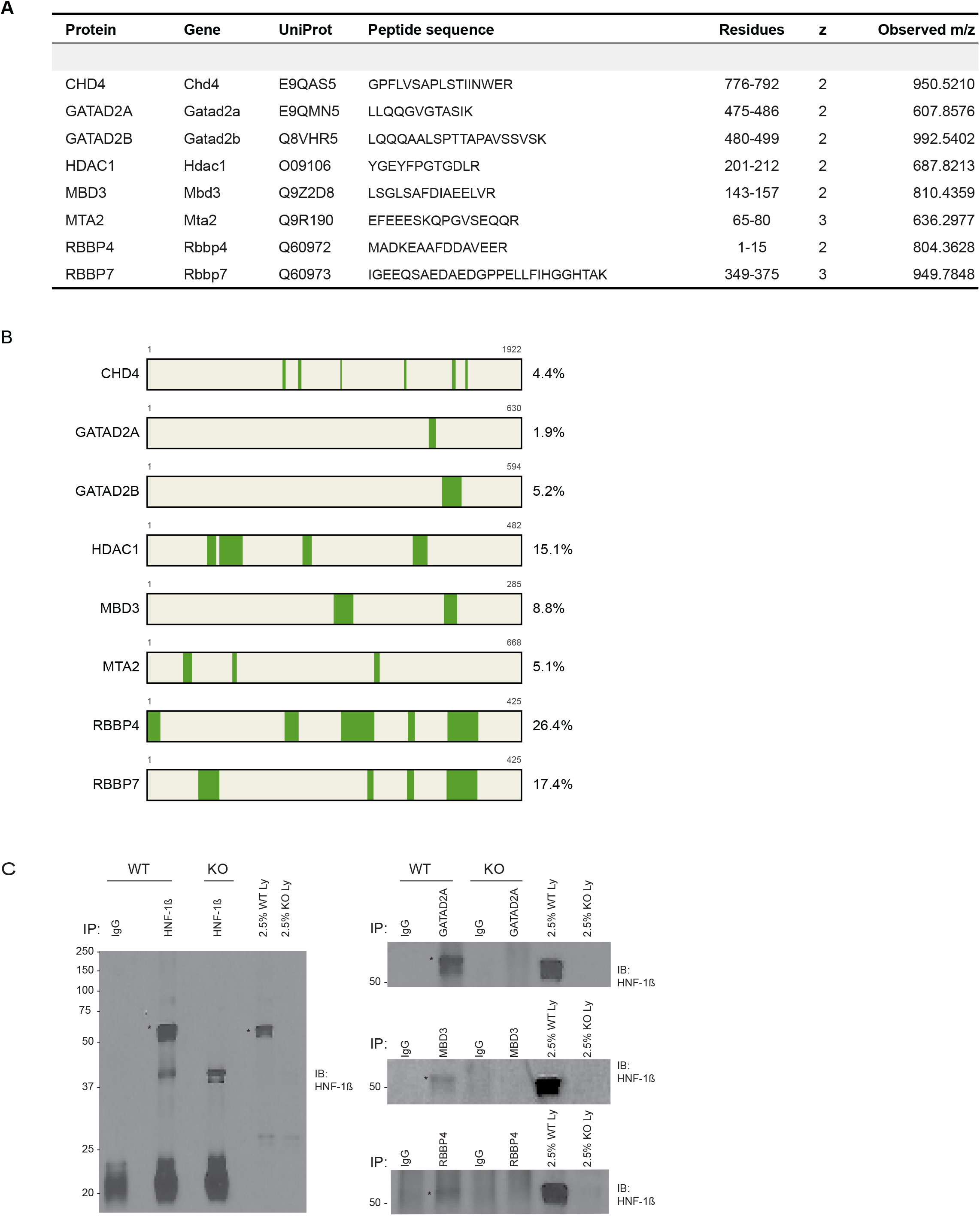
HNF-1β interacts with subunits of the nucleosome remodeling and deacetylase complex (NuRD). (A) RIME analysis of HNF-1β-interacting proteins in wild-type mIMCD3 cells identified multiple subunits of the NuRD complex. Peptide sequences, amino acid coordinates, and observed m/z are shown. (B) Coverage of peptides (green) derived from NuRD subunits CHD4, GATAD2A, GATAD2B, MBD3, MTA2, HDAC1, RBBP4 and RBBP7, which were identified by RIME in wild-type mIMCD3 cells. (C) Left: Western blot showing immunoprecipitation of HNF-1β using a rabbit polyclonal antibody in wild-type mIMCD3 cells (WT) but not in HNF-1β-deficient cells (KO). Immunoprecipitation with IgG was used as a negative control. * Indicates the position of HNF-1β. Molecular weight standards are shown on the left. Right: Western blots showing coprecipitation of HNF-1β following immunoprecipitation of NuRD subunits, GATAD2A (upper), MBD3 (middle), and RBBP4 (bottom) in wild-type mIMCD3 cells (WT) but not in HNF-1β-deficient cells (KO).

To further validate the protein-protein interactions and identify genomic sites of HNF-1β-NuRD interaction, we performed ChIP-seq on chromatin from WT mIMCD3 cells using polyclonal antibodies against three core NuRD subunits, GATAD2A, CHD4, and MBD3. The sites of NuRD subunit occupancy were compared with genomic sites of HNF-1β binding, which were also determined by ChIP-seq. We identified 27,627 total genomic sites that were occupied by GATAD2A, 32,111 sites that were occupied by CHD4, and 40,141 sites that were occupied by MBD3 (Fig.S3). Comparison with HNF-1β ChIP-seq revealed that 2963/27627 (10.7%) of GATAD2A-occupied sites, 2631/32111 (8.2%) of CHD4-occupied sites, and 2511/40141 (6.3%) of MBD3-occupied sites overlapped with HNF-1β binding sites. Conversely, 16.8%, 14.9%, and 14.2% of HNF-1β binding sites overlapped with GATAD2A-, CHD4-, and MBD3-occupied sites, respectively. Because a similar proportion of GATAD2A-, CHD4-, and MBD3-occupied sites exhibited colocalization with HNF-1β, we reasoned that genomic regions co-occupied by all three factors represent bona fide NuRD binding sites. Intersection of the three ChIP-seq datasets identified 18,469 high-confidence NuRD peaks in WT mIMCD3 cells of which 2776 peaks (15.0%) co-localized with HNF-1β (Fig. 3A-3B). Notably, 18.4% of the NuRD peaks in WT mIMCD3 cells colocalized with HNF-1β binding sites. Conversely, 2776 (15.8%) of the 17587 HNF-1β binding sites in WT mIMCD3 cells overlapped with NuRD-occupied peaks. This genomic colocalization of HNF-1β and the NuRD complex confirms the protein-protein interactions identified by RIME and suggests that their interaction could play a role in gene regulation.

**Figure 3.**
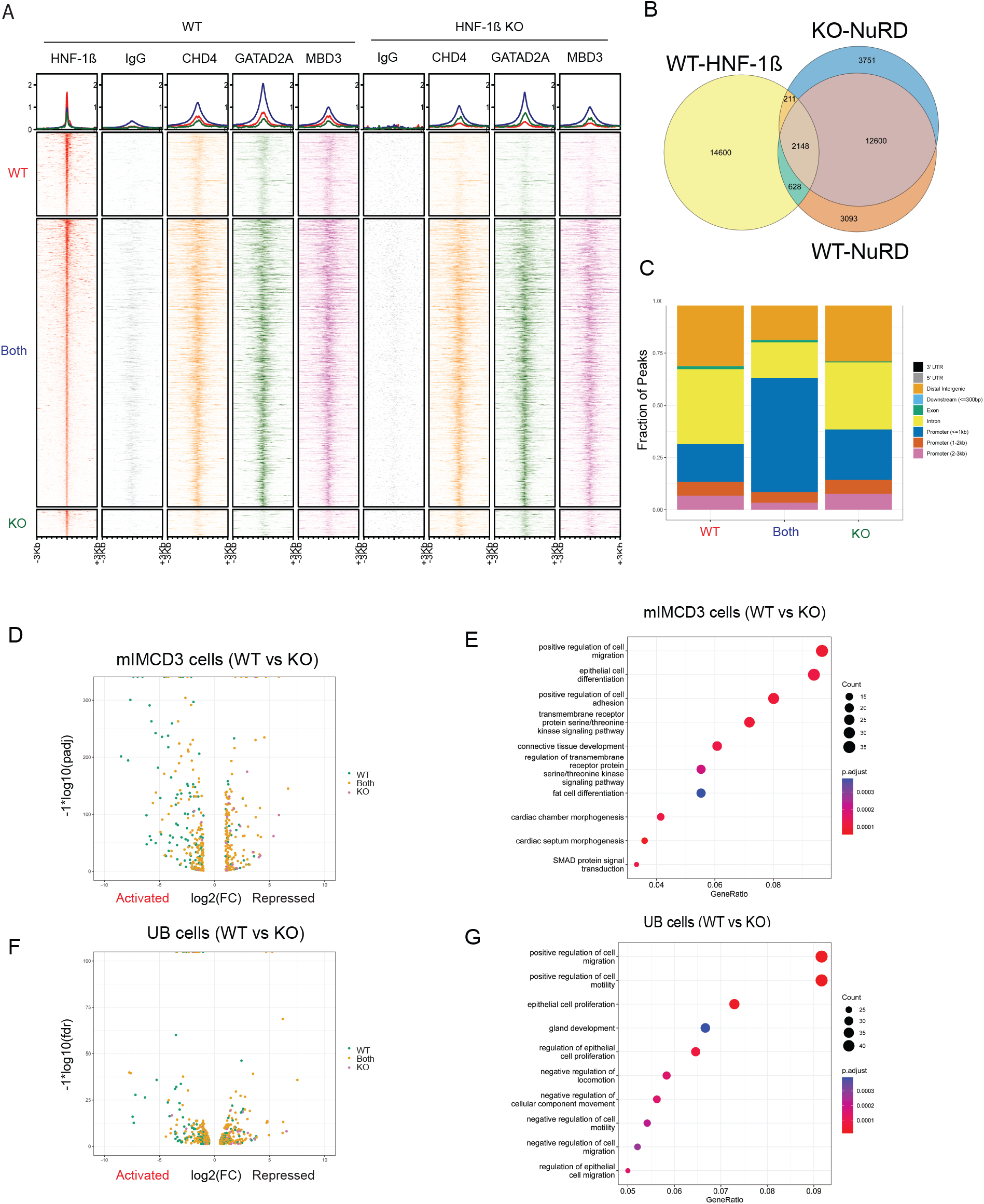
NuRD and HNF-1β are colocalized in the genome and regulate gene expression. (A) Heatmaps of ChIP-seq signals showing genomic colocalization of NuRD subunits CHD4, GATAD2A, and MBD3 centered at HNF-1β binding sites in wild-type (WT) and HNF-1β-deficient (KO) mIMCD3 cells. IgG was used as a negative control. Panel WT: shared NuRD and HNF-1β co-occupancy only in wild-type mIMCD3 cells. Panel Both: NuRD localization at HNF-1β binding sites in the presence and absence of HNF-1β. Panel KO: NuRD binding that was only detected in HNF-1β-deficient cells. Top panel shows aggregate ChIP-seq signals. (B) Venn diagrams show the overlap between HNF-1β and NuRD binding sites in wild-type (WT) and HNF-1β-deficient (KO) mIMCD3 cells. (C) Genomic distribution of unique and shared NuRD binding sites in the above three clusters. HNF-1β-dependent peaks (WT) are primarily located in introns (yellow) and distal intergenic regions (orange). (D) Volcano plot of genes that are differentially expressed in wild-type (WT) and HNF-1β-deficient (KO) mIMCD3 cells and contain a nearby colocalized HNF-1β and MBD3 binding site. Genes near a colocalized binding site that is only present in WT cells (green), in both WT and KO cells (gold), and only in KO cells (pink) are shown. (E) GO pathway analysis of differentially expressed genes. Dot size indicates the number of genes, and color indicates the adjusted p-value. (F) Volcano plot of genes that contain a nearby colocalized HNF-1β and NuRD binding site and are differentially expressed in ureteric bud (UB) cells from E14.5 wild-type mouse metanephroi (WT) compared to HNF-1β mutant metanephroi (KO). (G) GO pathway analysis of differentially expressed genes in HNF-1β mutant UB cells compared to wild-type UB cells.

### HNF-1β-NuRD complex regulates transcriptional activation in vitro in cultured kidney epithelial cells and in vivo in ureteric bud cells

Comparison of ChIP-seq results in WT and KO mIMCD3 cells revealed 628 HNF-1β binding sites (23% of the total WT sites) in which NuRD co-occupany was observed in WT cells but was absent or greatly reduced in KO cells, suggesting that HNF-1β was required to recruit NuRD to these sites (Fig. 3A-B). At 211 HNF-1β binding sites, NuRD occupancy was only observed in KO cells. Since NuRD itself lacks sequence-specific DNA binding capability, NuRD occupancy at these latter sites presumably reflects its interaction with alternative transcription factors that occupy HNF-1β binding sites in the absence of HNF-1β. The 628 unique sites in which NuRD occupancy is dependent on HNF-1β are primarily located in intronic and intergenic regions (Fig. 3C). This distribution suggests that NuRD genomic occupancy occurs mainly at distal regulatory elements, including putative enhancers, that may regulate HNF-1β-dependent transcription. To explore this possibility, we interrogated our previous transcriptomic datasets to identify HNF-1β-regulated genes that are differentially expressed in WT and KO cells and contain a nearby colocalized HNF-1β and NuRD site. A volcano plot (Fig. 3D) shows that NuRD-occupied sites that are dependent on the presence of HNF-1β (green) are located near genes that show higher expression in WT cells compared to KO cells, indicating that they are activated by HNF-1β (Table S2). These results suggest that HNF-1β-dependent recruitment of the NuRD complex is required for gene activation. To identify cellular pathways that are regulated by colocalized HNF-1β and NuRD peaks, we performed GO pathway analysis of the differentially expressed genes. We found enrichment of several pathways that are known to be regulated by HNF-1β, including cell migration and cell adhesion (Fig.3E).

To determine whether the changes in gene expression observed in cultured mIMCD3 cells are also seen in vivo, we interrogated the transcriptional profiles of ureteric bud (UB) cells purified from embryonic day 14.5 wild-type and HNF-1β mutant kidneys (22). We identified HNF-1β-regulated genes that are differentially expressed in WT and mutant UB cells and are located near a colocalized HNF-1β and NuRD-occupied site. A volcano plot shows that more genes that are down-regulated in UB cells from WT kidneys compared to HNF-1β mutant kidneys are located near a colocalized HNF-1β and NuRD binding site (Fig. 3F, Table S3). Pathway analysis of differentially expressed genes demonstrated enrichment of epithelial cell movement and cell differentiation (Fig.3G). Collectively, these results suggest that HNF-1β recruits NuRD to activate gene transcription both *in vitro* in cultured kidney epithelial cells and *in vivo* in the ureteric bud.

### HNF-1β-NuRD complex activates chromatin opening and histone H3K27 acetylation at distal enhancers

To understand the mechanisms whereby the HNF-1β-NuRD complex regulates gene expression, we performed ATAC-seq to measure genome-wide chromatin accessibility in WT mIMCD3 cells and KO cells (23, 24). A total of 20,273 ATAC-seq peaks were found in WT mIMCD3 cells, whereas 16,183 ATAC-seq peaks were detected in KO cells (Fig. S4A-B). Chromatin accessibility increased 25.3% in WT mIMCD3 cells compared to KO cells. We found a ∼5-fold increase of 2410 unique ATAC-seq peaks colocalized with HNF-1β binding sites in WT mIMCD3 cells compared to 480 ATAC-seq peaks in KO cells, suggesting that HNF-1β may open chromatin. Colocalized HNF-1β and ATAC-seq peaks were enriched in intergenic and intronic regions, suggesting that these colocalized peaks may function as active enhancers. We then compared the co-localized ATAC-seq peaks and HNF-1β binding sites with gene expression profiles in WT and KO mIMCD3 cells and UB cells. Co-localized ATAC-seq signals and HNF-1β binding sites were associated with HNF-1β-dependent transcriptional activation in both cell types (Fig. S4D-F). Pathway analysis of the differentially expressed genes with nearby colocalized ATAC-seq peaks and HNF-1β binding sites demonstrated enrichment of pathways involved in cell migration, chemotaxis, and axonogenesis (Fig. S4E-G).

Acetylation of lysine-27 on histone H3 (H3K27ac) is a hallmark of active enhancers and promoters that is strongly associated with transcriptionally active chromatin and distinguishes active enhancers from poised or inactive regulatory elements. CUT&RUN sequencing of H3K27ac sites in chromatin from WT mIMCD3 cells identified 6,201 H3K27ac sites that colocalized with HNF-1β genomic binding sites (Fig. S5A). 2,111 colocalized H3K27ac and HNF-1β sites were uniquely present in WT mIMCD3 cells, which is ∼4-fold higher than the 588 unique H3K27ac sites found in KO cells (Fig. S5B). Most of the colocalized HNF-1β and H3K27ac sites were located at putative enhancers that regulate both gene activation and repression in mIMCD3 cells and UB cells (Fig. S5C-F). Pathway analysis of the differentially expressed genes showed enrichment in cell migration, chemotaxis, and epithelial cell differentiation, pathways that are known to be regulated by HNF-1β (25). These results suggested that binding of HNF-1β at distal enhancers is associated with H3K27ac modifications.

Next, we examined H3K27ac modification and chromatin accessibility at the 628 unique genomic sites that exhibited HNF-1β-dependent co-occupancy by NuRD. We performed k-means clustering of the shared peaks and identified 113 unique chromatin-accessible sites that were co-occupied by HNF-1β and the NuRD complex and were enriched for H3K27ac in WT mIMCD3 cells (Fig 4A). In KO cells, these regions exhibited reduced chromatin accessibility and decreased H3K27ac. Most of the sites were located in introns or intergenic regions consistent with enhancers (Fig. 4B). Comparison with the RNA-seq transcriptomes of WT mIMCD3 cells and KO cells showed that the 113 co-occupied sites were located near genes that were down-regulated in KO cells compared to WT cells, indicating that they were activated by HNF-1β (Fig. 4C). We confirmed these findings *in vivo* by measuring the transcriptional profiles of ureteric bud (UB) cells purified from wild-type and HNF-1β mutant E14.5 metanephroi. Hierarchical clustering of RNA-seq datasets identified 54 differentially regulated genes including several well-characterized HNF-1β-regulated genes such as *Cdh16*, *Kif12*, *Notch1*, and *Dcdc2a* (Fig. 4C). IGV visualization of ChIP-seq datasets for HNF-1β and the NuRD complex subunits GATAD2A, CHD4, and MBD3, together with ATAC-seq and CUT&RUN analyses of H3K27ac and H3K4me3, demonstrated extensive colocalization of HNF-1β and NuRD at distal regulatory regions within the *Notch1*, *Arhgef16*, *Dcdc2a*, and *Bmp7* loci in WT mIMCD3 cells (Fig. S6). These sites exhibited features of active enhancers, including increased chromatin accessibility and enrichment of H3K27ac, supporting a role for HNF-1β-mediated recruitment of the NuRD complex to enhancer elements. Similarly, intronic enhancers are found in the loci of previously well characterized HNF-1β-dependent gene targets such as *Cdh16*, *Kif12* and *Pcsk9* in WT mIMCD3 cells (Fig.S7). Collectively, these findings support a new gene activation model whereby HNF-1β recruitment of the NuRD complex is associated with H3K27ac modification and opening chromatin.

**Figure 4.**
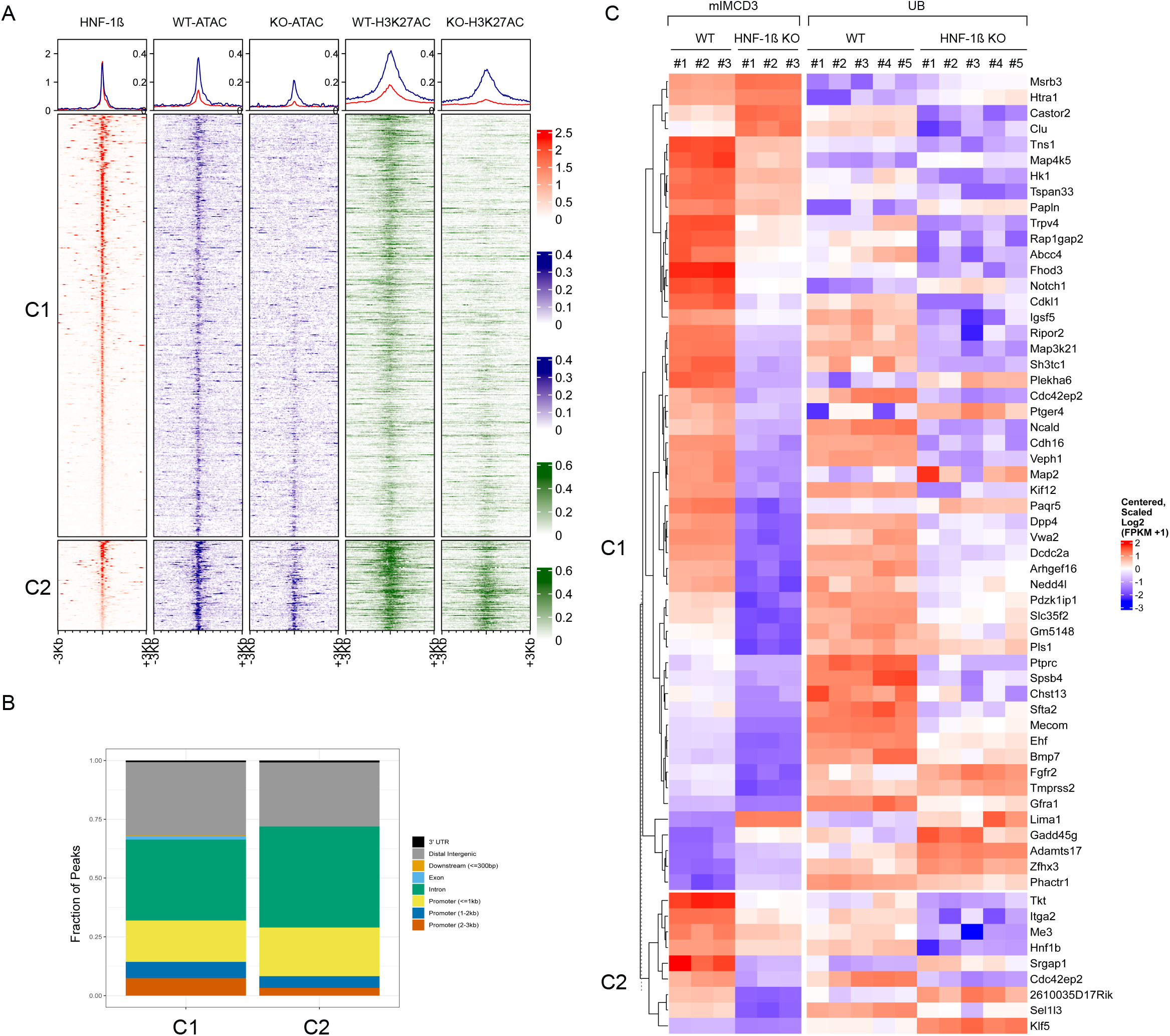
Colocalization of HNF-1β and NuRD is associated with increased chromatin accessibility and histone H3K27 acetylation. (A) Heatmaps show unsupervised (k-means) clustering of ATAC-seq and histone H3K27 acetylation (H3K27ac) centered at colocalized HNF-1β and NuRD binding sites in wild-type (WT) and HNF-1β-deficient (KO) mIMCD3 cells. Unsupervised clustering identified two distinct clusters, Cluster 1 (C1) and Cluster 2 (C2), which showed enrichment in WT cells compared to KO cells. Top panel shows aggregate signals of the HNF-1β ChIP-seq, ATAC-seq and H3K27ac CUT&RUN in C1 (red) and C2 (blue) in WT cells compared to KO cells. (B) Distribution of the genomic location of peaks identified in Cluster 1 and Cluster 2. (C) Hierarchical clustering shows differential expression of genes in Cluster 1 and Cluster 2 in WT mIMCD3 and UB cells compared to KO cells.

### NuRD subunit CHD4 is required for activation and repression of some HNF-1β-dependent genes

To test the functional importance of the interaction between HNF-1β and the NuRD complex, we inhibited the expression of NuRD core subunits CHD4 and GATAD2A and measured the effects on expression of HNF-1β target genes. Lentiviral-mediated expression of shRNA was used to knock down *Gatad2a* and *Chd4* in WT mIMCD3 cells (Fig 5A). We then measured the effects on global gene expression by RNA-seq (Fig. 5B). Transcriptomic profiling showed no significant difference between *Gatad2a* knockdown cells and control cells despite >2-fold knockdown of *Gatad2a*. This result may be due to GATAD2B, a homologous protein that may compensate for the loss of GATAD2A in kidney epithelial cells. In contrast, knockdown of *Chd4* in WT mIMCD3 cells resulted in the upregulation of 1,559 genes and the downregulation of 787 genes. Importantly, we identified a 67-gene signature associated with nearby HNF-1β and CHD4 co-occupancy, indicating that these genes are likely direct targets of HNF-1β-NuRD-mediated transcriptional regulation (Fig 5B). Gene ontology analysis demonstrated that these genes are associated with cell migration, chemotaxis, and axonal guidance (Fig.5C). Genome browser tracks at two representative loci, *Veph1* and *Slc6a19*, revealed colocalized HNF-1β and CHD4 occupancy at putative enhancer elements (Fig. 5D-E). Notably, CHD4 occupancy at these sites was absent in KO cells, indicating that recruitment of the NuRD complex to these enhancers is dependent on HNF-1β. These results indicate that CHD4 associated with the NuRD complex in HNF-1β occupied enhancers governs the expression of a subset of HNF-1β-dependent genes in kidney epithelial cells.

**Figure 5.**
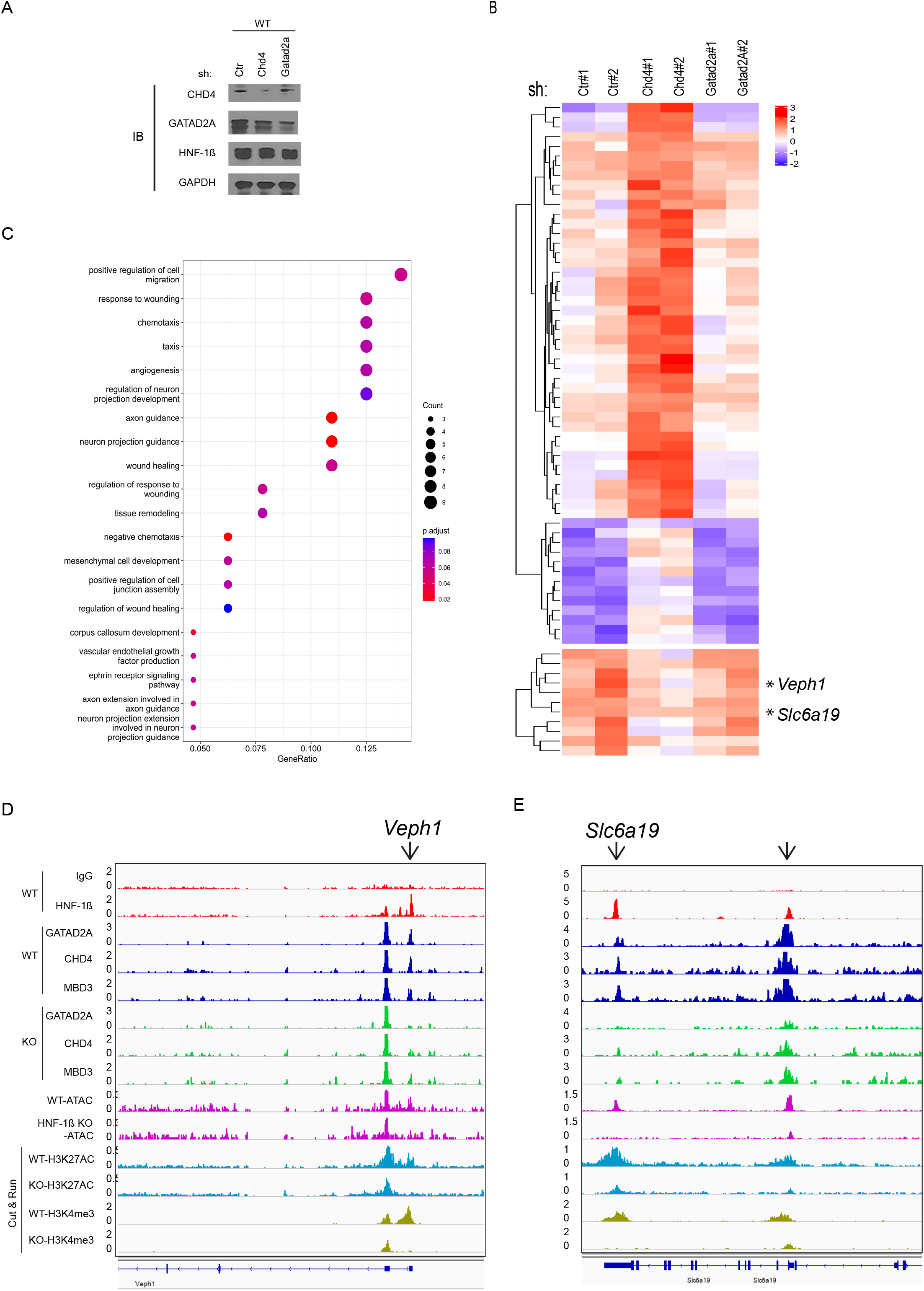
CHD4 is required for HNF-1β-dependent gene activation and repression. (A) Western blot showing protein levels of CHD4, GATAD2A, HNF-1β, and GAPDH (loading control) in WT mIMCD3 cells transduced with lentivirus expressing control shRNA (shCtr), shRNA targeting *Chd4* (sh*Chd4*), and shRNA targeting *Gatad2a* (sh*Gatad2a*). (B) Hierarchical clustering showing expression of 67 genes that are regulated by HNF-1β and contain a nearby colocalized HNF-1β and CHD4 binding site following knockdown of *Chd4* or *Gatad2a* in wild-type mIMCD3 cells. Up-regulated genes are shown in the upper two panels while down-regulated genes are shown in the bottom panel. (C) GO pathway analysis of the 67 gene set containing colocalized HNF-1β and CHD4 binding sites. (D-E) IGV gene tracks at *Veph1* (D) and *Slc6a19* (E) showing ChIP-seq of HNF-1β, GATAD2A, CHD4, and MBD3; ATAC-seq; and CUT&RUN sequencing of H3K27ac and H3K4me3 in wild-type (WT) and HNF-1β-deficient (KO) mIMCD3 cells. Arrows indicate the positions of enhancers.

## Discussion

Emerging evidence indicates that HNF-1β exerts long-range transcriptional control through cis-regulatory elements, highlighting an important role for distal enhancer-mediated gene regulation in maintaining epithelial homeostasis. Recently, we uncovered widespread reciprocal binding between HNF-1β and β-catenin that leads to hyperactivation of Wnt target genes in HNF-1β-deficient cells (26). This novel mechanism of HNF-1β in Wnt repression is due to direct competition between HNF-1β and β-catenin/LEF1 for binding to Wnt Responsive Elements (WREs). In addition, we demonstrated that HNF-1β has a role in downregulating β-catenin occupancy at WREs that are located more than 30 kb from target genes (27). These findings suggest that HNF-1β plays a role in regulating distal enhancers through chromatin remodeling or DNA looping mechanisms. However, the precise molecular mechanisms underlying HNF-1β-dependent long-range transcriptional regulation remain poorly understood.

Here, we employed an unbiased proteomic approach (RIME) to identify proteins that interact with endogenous HNF-1β in native chromatin. We identified several previously reported HNF-1β–interacting proteins, including PCBD1, Zyxin, and KPNA2 (18–20). The recovery of these known interaction partners supports the robustness and reproducibility of our RIME pipeline and validates its utility for identifying novel HNF-1β-associated proteins. The novel HNF-1β interactome that we identified contained multiple chromatin regulatory proteins and transcriptional cofactors (Fig.1B). The interacting proteins included components of the SWI/SNF chromatin remodeling complex (SMARCA5 and SMARCC2), the TIP60/NuA4 complex (RUVBL1 and RUVBL2), histone-modifying proteins (BAZ1B, TRIM28, CBX1, CBX3, and CBX5), transcription factors (GTF2I, HDGF, and TP53), and transcription elongation factors (SUPT16H, SUPT5H, and SUPT6H). Core subunits of the NuRD complex, including CHD4, GATAD2A, GATAD2B, MBD3, HDAC1, MTA2, RBBP4, and RBBP7, were among the most prominent proteins identified, suggesting that HNF-1β may regulate gene expression through interactions with multiple chromatin remodeling and transcriptional regulatory complexes.

We focused on the interaction of HNF-1β with the NuRD complex, because NuRD is a ubiquitous chromatin remodeling complex that plays important roles in cellular differentiation, embryonic stem cell biology, kidney development, and cancer (28). NuRD comprises modular subcomplexes containing interchangeable protein subunits that collectively regulate nucleosome compaction, nucleosome sliding, and chromatin accessibility through repositioning and eviction of histone octamers (29). The NuRD complex is highly evolutionarily conserved and consists of six core subunits organized in two subcomplexes (31). In mammals, NuRD comprises the ATP-dependent chromatin remodelers CHD3, CHD4, and CHD5; histone deacetylases HDAC1 and HDAC2; scaffold proteins MTA1, MTA2, and MTA3; methyl-CpG binding domain-containing proteins MBD2 and MBD3; histone-binding proteins RBBP4 and RBBP7; and GATAD2A and GATAD2B, which bridge the MBD and CHD subcomplexes to facilitate NuRD assembly and function. Structural studies indicate that the canonical NuRD complex contains one CHD, one MBD, one GATAD2, two HDAC, two MTA, and four RBBP subunits (32). Representatives of all six core subunits were identified as HNF-1β-interacting proteins. Since RIME was performed in cross-linked cells, the interactions between HNF-1β and individual NuRD subunits may be direct or indirect through common binding partners.

The NuRD complex was originally described as a transcriptional corepressor due to its intrinsic histone deacetylase activity and its widespread role in mediating transcriptional repression (30, 31). However, more recent studies suggest that it can also function as a coactivator in some situations (32). Paralog switching of the NuRD complexes may account for the functional differences in different cell types and at distinct genomic loci (32, 33). Several core components such as CHD3/4/5, GATAD2A/B, MBD2/3 and RBBP4/7 can interact with histones or methyl-CpG-rich sequences, which may facilitate NuRD’s genomic localization (34). However, NuRD lacks sequence-specific DNA-binding capability and is thought to be recruited to enhancers by its interaction with other sequence-specific DNA-binding transcription factors. Here, we found unexpectedly widespread co-localization of endogenous HNF-1β and NuRD in native chromatin from mouse renal epithelial cells. At ∼23% of co-occupied sites, NuRD occupancy was lost or greatly reduced in HNF-1β-deficient cells, indicating that HNF-1β was required for the recruitment of NuRD to the sites. Since NuRD in this study was defined stringently by the co-existence of MBD3, CHD4, and GATAD2A subunits, NuRD complexes that are composed of paralogous subunits were not included, and therefore the observed co-localization with HNF-1β is likely to be an underestimate of the true interaction. Closer examination of the NuRD and HNF-1β co-localized sites indicated that they were primarily located in introns and intergenic regions and were associated with increased chromatin accessibility and increased histone H3K27 acetylation, all characteristics of active enhancers. These results revealed a novel mechanism of transcriptional activation by HNF-1β through the recruitment of NuRD, histone H3K27 acetylation, and opening of chromatin. The interaction with NuRD may also be important for HNF-1β’s role as a pioneer transcription factor that remains bound to DNA during mitosis and rapidly restores HNF- 1β target genes after mitotic exit (35). Through sequence-independent tethering, NuRD has been shown to promote the binding of pioneer transcription factors to active enhancers and restrict binding to inappropriate genomic sites (36).

In addition to functioning as a transcriptional activator, HNF-1β can also suppress gene expression. For example, we have shown that HNF-1β inhibits Wnt signaling through transcriptional repression of *Lef1* (27). The C-terminal domain of HNF-1β has been shown to interact with the corepressor HDAC1, a histone deacetylase that increases the binding affinity of histones to DNA (7). Inhibition of deacetylase activity with trichostatin activates an HNF-1β reporter gene, consistent with a role in repression. Interestingly, HDAC1 as well as HDAC2 are core histone deacetylase components of NuRD (37) raising the possibility that the observed interaction with HDAC1 reflects a broader interaction with NuRD, similar to observations with other transcription factors such as GFI1 (1). Consistent with this possibility, sites of HNF-1β and

NuRD co-occupancy were also identified near genes that are upregulated in HNF-1β-deficient cells compared to wild-type cells, indicating that they are normally repressed by HNF-1β. NuRD represses transcription through a combination of ATP-dependent chromatin remodeling (mediated by CHD3, CHD4, or CHD5) and histone deacetylation (mediated by HDAC1 or HDAC2), which lead to chromatin compaction, restricted access by RNA polymerase, and silencing of target genes. For genes that are activated by the HNF-1β-NuRD complex, several mechanisms have been proposed whereby the normally repressive NuRD complex can function as a coactivator. In certain cellular contexts, the deacetylase activity of HDAC1 can itself be inhibited by protein acetylation thereby converting the NuRD complex from a corepressor to a coactivator (38). Methylation of the NuRD subunit MTA1 is required for NuRD’s repressive function, whereas demethylation of MTA1 switches it to a coactivator (39). The NuRD subunit CHD4 can function as a positive regulator of transcription by increasing super-enhancer accessibility (40). CHD4 has also been shown to maintain highly accessible chromatin at active enhancers while silencing cryptic enhancers (36). Further studies will be needed to determine whether these mechanisms or others underlie NuRD-dependent transcriptional activation by HNF-1β.

To confirm that the interaction between HNF-1β and NuRD is functionally significant, we knocked down the expression of the NuRD subunit CHD4 using shRNA in mIMCD3 cells that endogenously express HNF-1β. Transcriptional profiling of CHD4 knockdown cells compared to wild-type cells showed increased expression of most HNF-1β target genes that are located near sites of co-localized HNF-1β and NuRD. These findings are consistent with the well-established role of NuRD/CHD4 as a transcriptional repressor. In a smaller subset of genes, knockdown of CHD4 decreased the expression of nearby HNF-1β target genes. These findings indicate that the interaction between HNF-1β and NuRD is functionally important for transcriptional activation as well as repression. The relationship between HNF-1β and NuRD may help explain previous studies on the role of NuRD in kidney development (28). Knockout of CHD4 (also known as Mi-2β) in mice results in renal hypoplasia, a phenotype that is also seen in humans carrying mutations of HNF1B, suggesting that the CHD4 knockout phenotype could in part be due to an impairment in HNF-1β-dependent gene regulation (41). CHD4 has also been shown to interact with Sal1, another transcription factor that is essential for kidney development (42). Disruption of the interaction between Sal1 and CHD4 leads to defects in nephron patterning including absence of the loops of Henle (43). An identical phenotype is observed in *Hnf1b* knockout mice (44), suggesting that NuRD may play a common role. It is also noteworthy that the nephron patterning defects seen in *Hnf1b* and *Sal1* mutant mice have been attributed to defects in Notch signaling (43, 44), and *Notch1* was identified as one of the genes that is regulated by co-localized HNF-1β and NuRD in our studies (Figs. 4C, S6A).

In summary, our findings establish a new role of HNF-1β in genomic targeting of the NuRD complex to distal enhancers to activate gene expression. More broadly, the HNF-1β interactome that we identified by RIME includes a variety of chromatin remodeling complexes, histone modifiers, transcription factors, and elongation factors, which suggests that diverse mechanisms of action of HNF-1β-dependent gene regulation exist. Understanding how HNF-1β exerts transcriptional control through its protein-protein interactions and long-range chromatin remodeling functions could be highly relevant to embryonic kidney development and the pathogenesis of human *HNF1B* nephropathy.

## Methods

### Cell culture

mIMCD3 cells and 293T cells were obtained from ATCC and maintained in DMEM supplemented with 10% FBS and antibiotics (100 units/ml penicillin and 100 µg/ml streptomycin). HNF-1β-deficient mIMCD3 cells were generated as described previously (17). Cells were cultured at 37°C under 5% CO_2_.

### Rapid Immunoprecipitation Mass Spectrometry of Endogenous Proteins (RIME)

Wild-type mIMCD3 and HNF-1β-deficient cells were seeded in 10-cm culture dishes at a density of 5 × 10^5^ cells per dish and maintained in DMEM supplemented with 10% fetal bovine serum (FBS). Cells were cultured to 80–90% confluence, after that the medium was replaced with fresh DMEM containing 10% FBS and incubated for an additional 16 hrs before crosslinking. Cells were crosslinked with 1% paraformaldehyde for 10 mins at room temperature and processed for chromatin immunoprecipitation using SimpleChIP Plus Enzymatic Chromatin Kits (Cell Signaling #9005) according to the manufacturer’s instructions. HNF-1β-containing chromatin complexes were immunoprecipitated using a rabbit polyclonal anti-HNF-1β antibody (Santa Cruz Biotechnology, sc-22840). Parallel immunoprecipitations were performed using HNF-1β-deficient cells as a negative control for subsequent mass spectrometry analysis. Briefly, 5 μg of anti-HNF-1β antibody was used for each immunoprecipitation from the nuclear extract prepared from one-half of a 10-cm dish of confluent cells. Following the final wash, bead-bound immune complexes were washed twice with 1 mL of freshly prepared 100 mM ammonium bicarbonate (NH4HCO3, AMBIC) maintained at 4°C. The beads were then transferred to a fresh microcentrifuge tube and subjected to on-bead digestion with sequencing-grade trypsin (Promega). Twenty microliters of trypsin (10 ng/μL) were added, and samples were incubated overnight at 37°C. An additional 20 μL of trypsin (10 ng/μL) was subsequently added, and digestion continued for 4 hours at 37°C to ensure complete proteolysis. Following digestion, magnetic beads were separated using a magnetic rack, and the peptide-containing supernatant (∼40 μL) was collected. Peptides were acidified by adding neat trifluoracetic acid (TFA) to a final concentration of 5% (v/v). Post-digestion samples were dried in a vacuum centrifuge. The samples were reconstituted in 50mM triethylammonium bicarbonate (TEAB) with 4mM TCEP and incubated for 30 min at 37°C. Iodoacetamide in 50 mM TEAB was added to a final concentration of 8 mM and incubated in the dark at room temperature for 30 min. The sample was diluted 1:1 with 98% water, 2% acetonitrile, and 0.1% trifluoracetic acid (TFA). The pH was checked and adjusted to pH 3 with 10% TFA. The samples were cleaned with a C18-like Stage tip (45).

### Liquid Chromatography and Mass Spectrometry

We analyzed approximately 500 nanograms of peptide (reconstituted in 97.7:2:0.1, H2O: acetonitrile (ACN): formic acid (FA)) by capillary LC-MS with an Easy NanoLC 1100 chromatography system (Thermo Scientific, Waltham, MA) in-line with an Orbitrap Fusion mass spectrometer (Thermo Scientific) in data dependent acquisition mode. We separated peptides on a 40 cm, 100 micrometers internal diameter (ID) C18 column packed in-house with Dr Maisch 1.9 micrometer C18aq with 120 angstrom pore size; the column temperature was held at 55 °C. We eluted peptides at a flowrate of 300 nl/min over a linear gradient with the following profile: 5 – 22% solvent B from 0 – 45 minutes, 22 – 35% solvent B from 45 – 70 minutes, 35 – 90 % solvent B from 70 – 80 minutes, then held at 90% B until 85, where solvent A was 0.1% FA in water and solvent B was 0.1% FA in ACN. We employed a data dependent acquisition mode with the top 12 most abundant ions per cycle submitted for tandem MS mode. We employed the following MS parameters: ESI voltage 2.1 kV, ion transfer tube 275 °C; Orbitrap MS1 scan 60k resolution in profile mode from 380 – 1580 *m/z* with 250 msec maximum injection time; 4 × 10E5 automatic gain control (AGC); MS2 was triggered on precursors with 2 – 7 charges above 5000 counts; MIPS (monoisotopic peak determination) was set to Peptide; MS2 settings were: 1.6 Da quadrupole isolation window, MS2 triggered on precursors charge states 2 – 7 above 5000 counts; precursor isolation window 1.6 Da; MIPS (monoisotopic peak determination) set to Peptide; collision induced dissociation (CID) activation (35% collision energy, 10 msec activation time and 0.25 activation Q) and ion trap detection, 35 msec injection time, 1 × 10 E4 AGC, dynamic exclusion duration of 15 sec with +/-10 ppm mass tolerance. We processed peptide tandem MS using SEQUEST (Thermo Scientific) in Proteome Discoverer 3.0. The mouse Universal Proteome database (UP000000589) was downloaded from UniProt on 8/1/2022 and merged with a lab contaminant protein database (https://github.com/HaoGroup-ProtContLib). We applied the precursor mass recalibration node with precursor mass tolerance 20 ppm, product ion tolerance 0.8 Da with fixed carbamidomethyl (CAM) modification of cysteine 57.0215 *m/z*. The SEQUEST database search parameters included enzyme trypsin full specificity, 2 missed cleave sites peptide length 6 – 50 amino acids long; precursor tolerance 15 ppm, fragment ion tolerance 0.6 Da and maximum 4 dynamic modifications per peptide (46). We specified CAM cysteine (+57.021 Da) as a fixed modification and the dynamic modifications were acetylation of protein N-terminus (+42.011 Da), methionine loss at the protein N-terminus (-131.040 Da), methionine loss + acetylation at the protein N-terminus (-89.030 Da), oxidation of methionine (+15.995 Da), conversion of glutamine to pyroglutamic acid (-17.027 Da), and deamidation of asparagine or glutamine (+0.984 Da). We applied 1% protein and peptide False Discovery Rate (FDR) filters using the Percolator algorithm in Proteome Discoverer 3.0. (16).

### Chromatin immunoprecipitation and sequencing (ChIP-seq)

Wild-type mIMCD3 cells and HNF-1β-deficient cells were seeded in 10-cm tissue culture dishes (5×10^6^ cells/dish) and cultured in DMEM containing 10% FBS. Cells were grown to 80−90% confluence, after incubation for an additional 16 hrs, cells were fixed in 1% paraformaldehyde for 10 mins at room temperature. Chromatin immunoprecipitation was performed using an antibody against GATAD2A (Abcam, ab188472) CHD4 (Abcam, ab240640), MBD3 (Abcam, ab157464) or isotype-specific IgG (Cell Signaling #2729) as a negative control. ChIP was performed using SimpleChIP Plus Enzymatic Chromatin Kits (Cell Signaling #9005) according to the manufacturer’s instructions. For ChIP-seq, ∼100 pg of eluted DNA or input was used for library creation with ThruPLEX DNA-seq Kits (Takara Bio). Sequencing was performed by the University of Minnesota Genomics Center using an Illumina Next-seq P2 in 1×100bp single-read mode.

### ATAC-sequencing

Wild-type mIMCD3 cells and HNF-1β-deficient cells were seeded in triplicate in 6-well plates (5×10^5^ cells/well) and cultured in DMEM containing 10% FBS. Cells were grown to 80−90% confluence and changed with fresh DMEM containing 10% FBS. After incubation for an additional 16 hrs, 50,000 cells were isolated after trypsin digestion and were washed once with iced cold 200µl 1XPBS and resuspended in 100 μl of cold lysis buffer (10 mM Tris-HCl pH 7.4, 10 mM NaCl, 3 mM MgCl2, 0.1% IGEPAL CA-630), spun at 500 g for 10 min at 4 ^O^C and resuspended in in 50 μl of the transposition reaction mix. Transposition occurred at 37°C for 30 min, after which transposed, DNA was purified using a Qiagen Min Elute Kit and eluted in 10 μl Elution Buffer. ATAC-Seq libraries were generated using dual-indexed Nextera reagent kits by using transposed DNA fragments according to the manufacturer’s instructions. All libraries were combined into a single pool. Sequenced the pool across in a single lane of a HiSeq 2500 HO 2×50-bp run using v4 chemistry.

### CUT&RUN sequencing

Wild-type mIMCD3 cells and HNF-1β-deficient cells were seeded in 24-well tissue culture plate (5×10^4^ cells/well) and cultured in DMEM containing 10% FBS. Cells were grown to 80−90% confluence, after incubation for an additional 16 hrs, cells were trypsin digested and 250,000 cells were used for each CUR&RUN reaction. Chromatin immunoprecipitation was performed using an antibody against H3K27ac (Cell Signaling #8173) H3K4me3 (Cell Signaling #9751) or isotype-specific IgG (Cell Signaling #2729) as a negative control. CUT&RUN was performed using CUT&RUN kit (Cell Signaling #86652) according to the manufacturer’s instructions. For CUT&RUN, ∼100 pg of eluted DNA was used for library creation with ThruPLEX DNA-seq Kits (Takara Bio). Sequencing was performed by the University of Minnesota Genomics Center using an Illumina Next-seq 2000 in 2×50bp paired-end mode.

### RNA-sequencing

Briefly, wild-type mIMCD3 cells and HNF-1β-deficient cells were seeded in triplicate in 6-well plates (5×10^5^ cells/well) and cultured in DMEM containing 10% FBS.

Cells were grown to 80−90% confluence and infected with lentivirus shRNA against *Gatad2a* and *Chd4*. Lentivirus generation was performed as described previously (47). Targeting DNA sequences used for this study are listed in Table S4. After 72 hrs. lentiviral post-induction, cells were treated with 100 ng/ml Wnt3a (R&D Systems) or vehicle alone as a negative control. After incubation for an additional 16 hrs, RNA was harvested using RNeasy Mini Kits (Qiagen). TruSeq stranded mRNA libraries were created from each sample and sequenced in a 50 bp PE run on a NovaSeq instrument the University of Minnesota Genomics Center.

### Co-immunoprecipitation and immunoblot

Wild-type mIMCD3 and HNF-1β-deficient cells were seeded in 10-cm culture dishes at a density of 5 × 10^5^ cells per dish and maintained in DMEM supplemented with 10% fetal bovine serum (FBS). Cells were cultured to 80–90% confluence, after that the medium was replaced with fresh DMEM containing 10% FBS and incubated for an additional 16 hrs before crosslinking. Cells were crosslinked with 1% paraformaldehyde for 10 mins at room temperature and processed for chromatin immunoprecipitation using SimpleChIP Plus Enzymatic Chromatin Kits (Cell Signaling #9005) according to the manufacturer’s instructions. For each co-immunoprecipitation (co-IP), 40% of the sonicated, crosslinked chromatin prepared from a single 10-cm dish was used. Chromatin-associated protein complexes were immunoprecipitated using antibodies against HNF-1β, GATAD2A, MBD3, or RBBP4, with normal rabbit IgG serving as a negative control. Parallel immunoprecipitations were performed using HNF-1β-deficient cells as a negative control. Instead of proceeding with the DNA elution step of the ChIP protocol, proteins bound to Protein-G magnetic beads were eluted by adding 30µl of 1X Laemmli protein sample buffer, followed by heating at 95 ^0^C for 10 mins. Half of each immunoprecipitate and 2.5% of the nuclear lysate (Input) were subjected to immunoblot blot analysis. Immunoblot analysis was performed as described previously (47). Antibodies used for the studies are listed in Table S5.

### ChIP-Seq Data Analysis

Low input, dual-indexed Rubicon Thruplex libraries were sequenced on an Illumina NextSeq 2000 in a 1×100bp configuration with a mean depth of 50 million. Quality assessment was performed using FastQC (0.11.7) https://www.bioinformatics.babraham.ac.uk/projects/fastqc/. Reads were trimmed with TrimGalore (v0.6.0) and mapped to GRCm38.p6 with bwa mem (v0.7.17-r1188) and sorted with samtools (v1.21) (48–50) . Peaks were identified using MACS (v2.1.1.20160309) using default parameters (51). Genomic regions were associated with genes using the ChIPseeker (1.32.0) R package defining promoters at 3 kb upstream or downstream of the transcription start site. Figures were generated using EnrichedHeatmap (1.26.0) and ggplot2(3.3.6) (52, 53).

### CUT&RUN Data Analysis

Low input, dual-indexed Rubicon Thruplex libraries were sequenced on an Illumina NextSeq 2000 in a 2×51 bp configuration with a mean depth of 21 million. Quality assessment was performed using FastQC (0.11.7). Reads were trimmed with TrimGalore (v0.6.0) and mapped to GRCm38.p6 with bwa mem (v0.7.17-r1188) and sorted with samtools (v1.21). Non-unique reads were removed with Picard Mark Duplicates (2.18.16-SNAPSHOT). Peaks were identified using MACS (v2.1.1.20160309) using the “-m 2 20 --keep-dup all -f BAMPE” options and the IgG samples as controls. Genomic regions were associated with genes using the ChIPseeker (1.32.0) R package defining promoters at 3kb upstream or downstream of the transcription start site. Figures were generated using EnrichedHeatmap (1.26.0) and ggplot2 (3.3.6).

### ATAC-seq Data Analysis

Dual-indexed Nextera libraries were sequenced on an Illumina NextSeq 2500 in a 2×50bp configuration. Quality assessment was performed using FastQC (0.11.7). Reads were trimmed with TrimGalore (v0.6.0) and mapped to GRCm38.p6 with bwa mem (v0.7.17-r1188) and sorted with samtools (v1.21). Non-unique reads were removed with Picard Mark Duplicates (2.18.16-SNAPSHOT, Broad Institute) resulting in 19 or 13 million unique reads for wild-type or HNF-1β KO cells respectively. Peaks were identified using MACS (v2.1.1.20160309) using the “--broad -q 0.05 –nomodel” options. Figures were generated using EnrichedHeatmap (1.26.0) and ggplot2 (3.3.6).

RNA-seq Data Analysis. RNA-seq libraries were sequenced on an Illumina Novaseq S4 in a 2×150 configuration with a mean read depth of 41 million. Quality assessment was performed using FastQC (0.11.7). Reads were trimmed with Trimmomatic (0.33) and mapped to GRCm38 using HiSat2 (2.1.0) (54). Gene level abundance was estimated using Rsubread package in R (3.5.0) and Differential expression analysis was performed using glmQLFit function in EdgeR (55, 56). Gene ontology analysis was performed with ClusterProfiler (v4.4.4) (57) .

### Statistical analysis

Student’s two-tailed unpaired t test was used for pairwise comparisons. P<0.05 was considered significant.

## Author Contributions

SC conducted experiments. MDG and ZD conducted ChIP-seq, RNA-seq and ATAC-seq and CUR&RUN analysis. LH and TM performed RIME analysis. AS provided UB RNA-seq dataset. SC and PI designed experiments, analyzed data, and wrote the manuscript.

## Acknowledgments and Financial Support

We thank Mark Murphy for advice on ATAC-seq, Illumina sequencing services performed by the University of Minnesota Genomics Center, and the Minnesota Supercomputing Institute provided bioinformatic, software support and data storage. Trinity Hausch for her artwork. Supported by NIH grant R01 DK42921 (P.I.) and R01 HD120295 (M.D.G). The Orbitrap Eclipse instrumentation platform used in this work was purchased through High-end Instrumentation Grant S10OD028717 from the NIH.

## Disclosures

None

## Notes

### Competing Interest Statement

The authors have declared no competing interest.

