## Supplement for "Hepatocyte nuclear factor-1β recruits the nucleosome remodeling and deacetylase complex to activate gene transcription"

**Supplemental Tables**

**Table S1. List of the HNF-1β interacting proteins identified by RIME**

**Table S2. List of differentially expressed genes associated with co-localized HNF- 1β and NuRD binding sites uniquely in wild-type mIMCD3 cells**

**Table S3. List of differentially expressed genes associated with co-localized HNF-1β and NuRD uniquely *in vivo* UB cells**

**Table S4. List of primers used for the construction of shRNA KD plasmids**

**Table S5. List of antibodies**

### Supplemental Figures

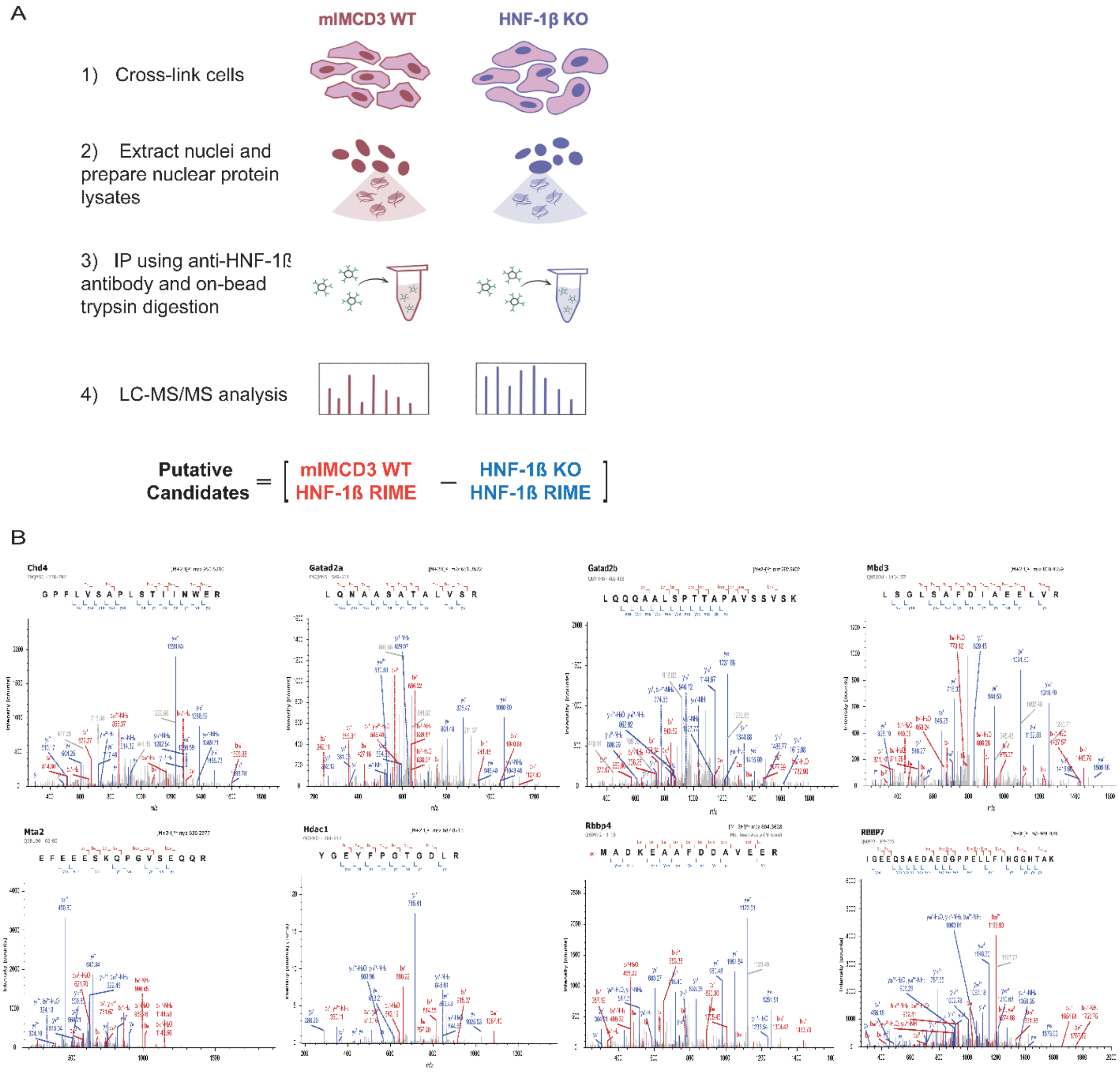


**Figure S1. *Identification of HNF-1β-interacting proteins by rapid immunoprecipitation mass spectrometry of endogenous proteins (RIME).*** (A) Schematic diagram showing the experimental approach. Wild-type mIMCD3 cells (WT) were cross-linked, HNF-1β was immunoprecipitated from nuclear lysates with a rabbit polyclonal antibody, coprecipitated proteins were digested on-bead, and generated peptides were identified by LC-MS/MS. RIME analysis of HNF-1β-deficient mIMCD3 cells (KO) was performed to exclude non-specific binding. (B) Tandem MS/MS spectra of doubly charged peptides corresponding to core NuRD subunits, CHD4, MBD3, GATAD2A, GATAD2B, MTA2, HDAC1, RBBP4 and RBBP7, which were identified by RIME in WT cells. Red represents b-fragment ions generated from the C-terminal sequence; blue represents y-fragment ions generated from the N-terminal sequence. Intensity (counts) is plotted against m/z.

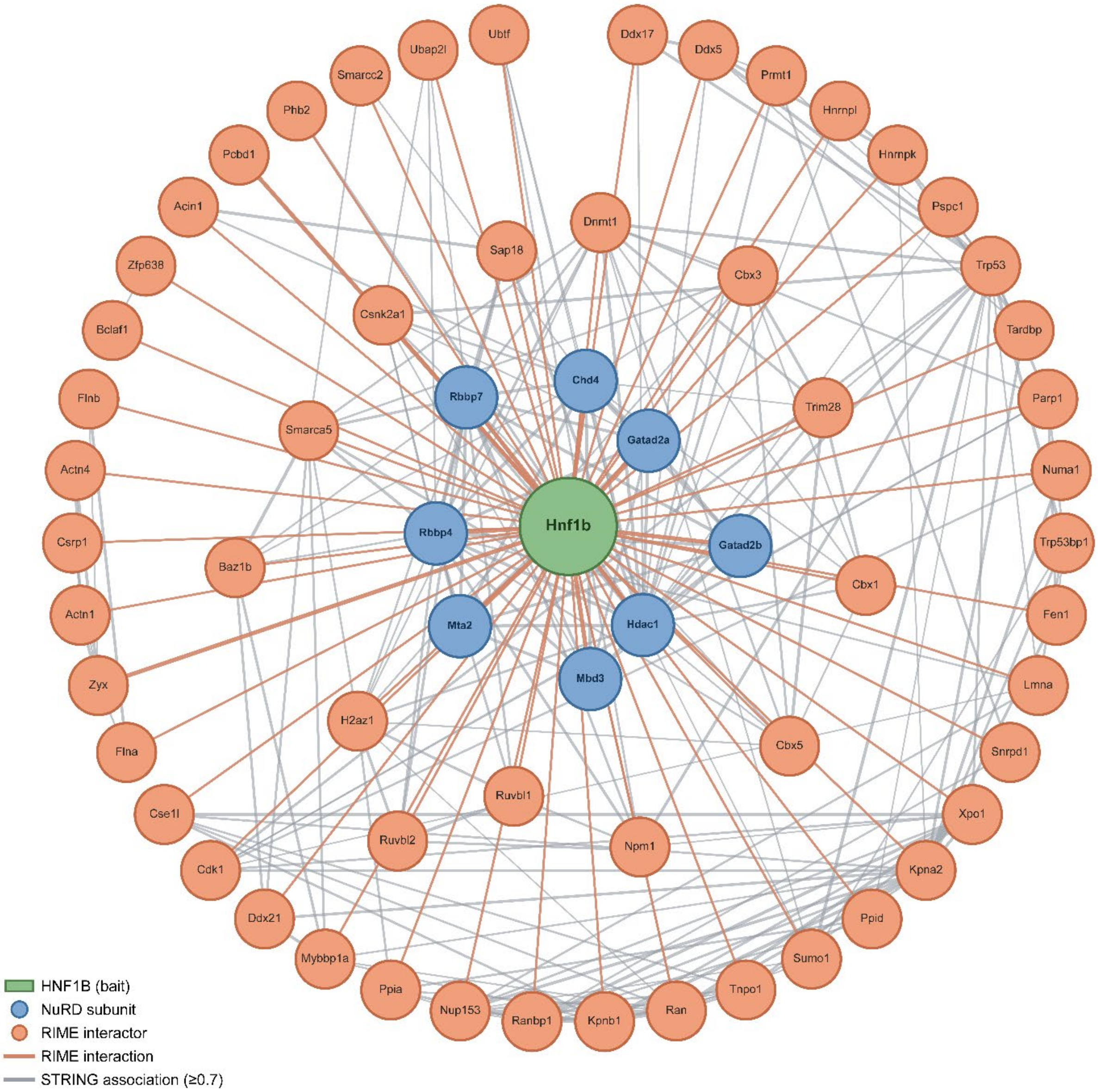


**Figure S2.** ***The interactome of HNF-1β and the NuRD complex.*** (A) 484 HNF-1β-interacting proteins identified with RIME were queried against the STRING database [v12.0; *Mus musculus*] using a minimum interaction score of 0.7 (high confidence). The resulting association network was imported into Cytoscape [v3.10.4]. Known HNF-1β-interacting proteins (Zyx, Pcbd1, and Kpna2) and NuRD subunits (Chd4, Gatad2a, Gatad2b, Hdac1, Mbd3, Mta2, Rbbp4, Rbbp7) were selected as seed nodes, and their first-neighbor subnetwork was extracted for visualization. The final network comprised 63 nodes — the Hnf1b bait, 8 NuRD subunits, and 54 first-neighbor RIME interactors. Nodes were colored by class (Hnf1b bait, NuRD subunit, RIME interactor). Edges denote high-confidence STRING associations (combined score ≥ 0.7; grey), and direct RIME interactions with Hnf1b are shown in orange.

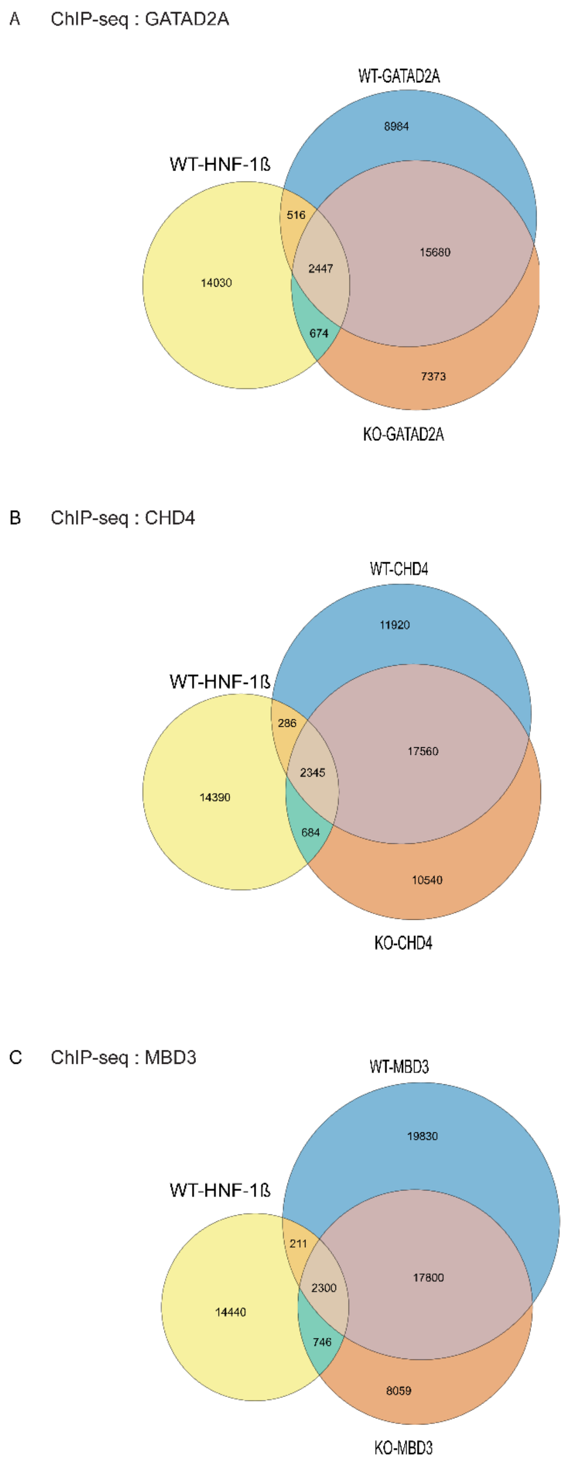


**Figure S3. ChIP-seq analysis of NuRD components, GATAD2A, CHD4 and MBD3.** Venn diagrams show the overlap between HNF-1β binding sites in wild-type (WT) mIMCD3 cells and genomic binding sites for NuRD subunits, GATAD2A (A), CHD4 (B), or MBD3 (C) in WT cells or HNF-1β-deficient (KO) cells.

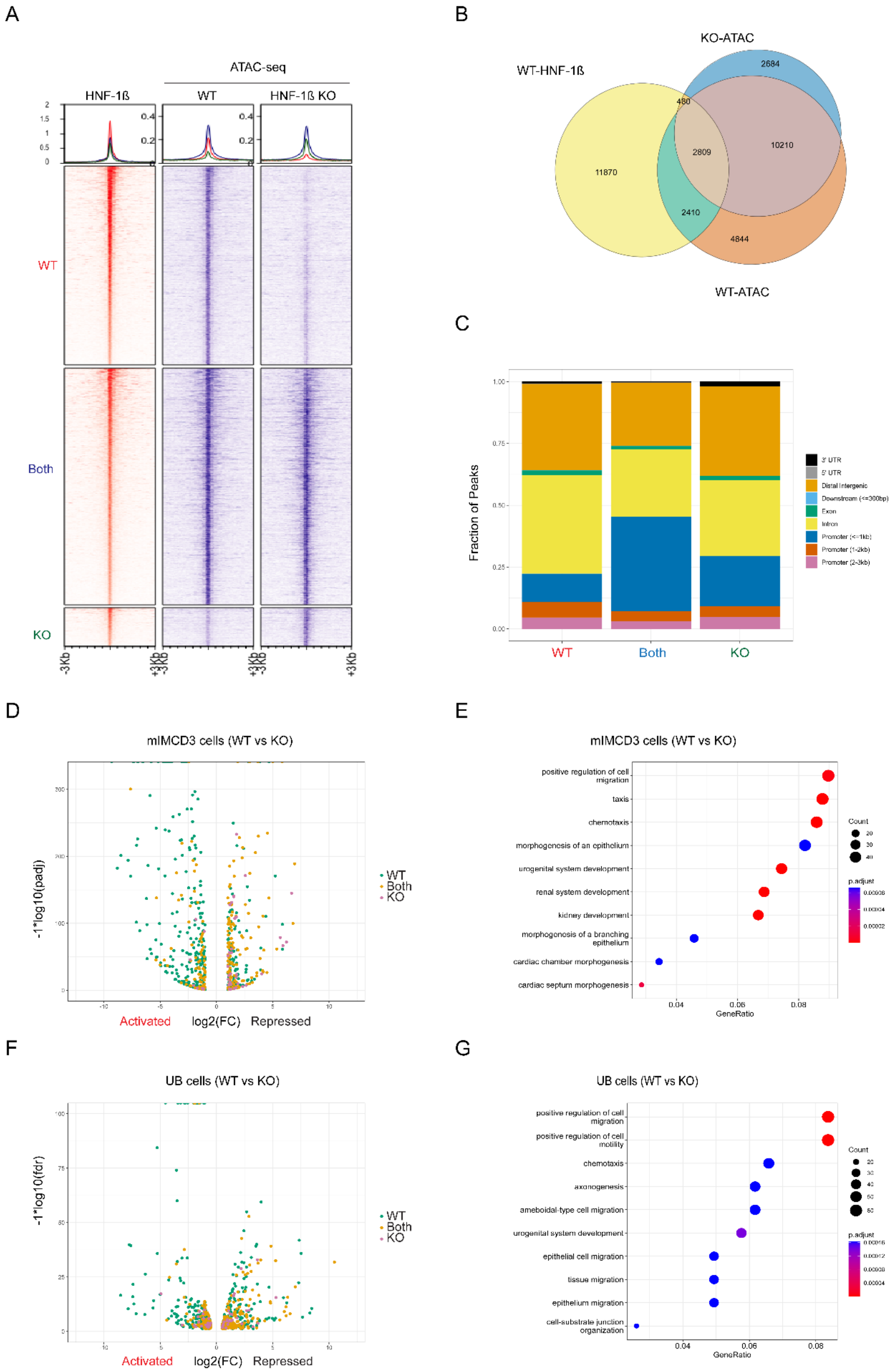
A
ATAC-seq
HNF-1ß
WT
HNF-1ß KO
WT
Both
KO
-3Kb
+3Kb
B
KO-ATAC
WT-HNF-1ß
WT-ATAC
2684
480
2809
10210
11870
2410
4844
C
Fraction of Peaks
3' UTR
5' UTR
Distal Intergenic
Downstream (<=300bp)
Exon
Intron
Promoter (<=1kb)
Promoter (1-2kb)
Promoter (2-3kb)
WT
Both
KO
D
mIMCD3 cells (WT vs KO)
-1*log10(padj)
WT
Both
KO
Activated
log2(FC)
Repressed
E
mIMCD3 cells (WT vs KO)
positive regulation of cell migration
taxis
chemotaxis
morphogenesis of an epithelium
urogenital system development
renal system development
kidney development
morphogenesis of a branching epithelium
cardiac chamber morphogenesis
cardiac septum morphogenesis
Count
p.adjust
GeneRatio
F
UB cells (WT vs KO)
-1*log10(fdr)
WT
Both
KO
Activated
log2(FC)
Repressed
G
UB cells (WT vs KO)
positive regulation of cell migration
positive regulation of cell motility
chemotaxis
axonogenesis
ameboidal-type cell migration
urogenital system development
epithelial cell migration
tissue migration
epithelium migration
cell-substrate junction organization
Count
p.adjust
GeneRatio

**Figure S4. HNF-1β regulates chromatin accessibility at distal enhancers.** (A) Heatmaps show ATAC-seq signals in wild-type mIMCD3 cells (WT) and HNF-1β-deficient cells (KO) centered on HNF-1β binding sites (left lane). Panel WT: Unique ATAC-seq signals colocalized with HNF-1β only in WT cells. Panel Both: Colocalized ATAC-seq signals at HNF-1β binding sites in the presence and absence of HNF-1β. Panel KO: Unique ATAC-seq signals in HNF-1β-deficient cells. Top panel shows the aggregate ATAC-seq signals. (B) Venn diagram shows the overlap between HNF-1β binding sites in WT cells and ATAC-seq signals in WT and KO cells. (C) Distribution of genomic locations of unique and shared ATAC-seq signals in the above three clusters (WT, Both and KO). HNF-1β-dependent peaks (WT) are primarily located in introns (yellow) and distal intergenic regions (orange). (D) Volcano plots of genes that contained nearby colocalized HNF-1β and ATAC-seq peaks and were differentially expressed in WT cells and KO cells. Genes with a nearby colocalized HNF-1β and ATAC-seq peak only in wild-type mIMCD3 cells (green), in both wild-type and HNF-1β-deficient cells (gold), and only in HNF-1β-deficient cells (pink) are shown. (E) GO pathway analysis of HNF-1β-regulated genes located near colocalized HNF-1β and ATAC-seq peaks. Dot size indicates the number of genes, and color indicates the adjusted p-value. (F) Volcano plots of genes with nearby HNF-1β and ATAC-seq peaks that are differentially expressed in ureteric bud (UB) cells from E14.5 wild-type mouse kidneys (WT) and HNF-1β mutant kidneys (KO). (G) GO pathway analysis of differentially expressed genes located near colocalized HNF-1β and ATAC-seq peaks in UB cells.

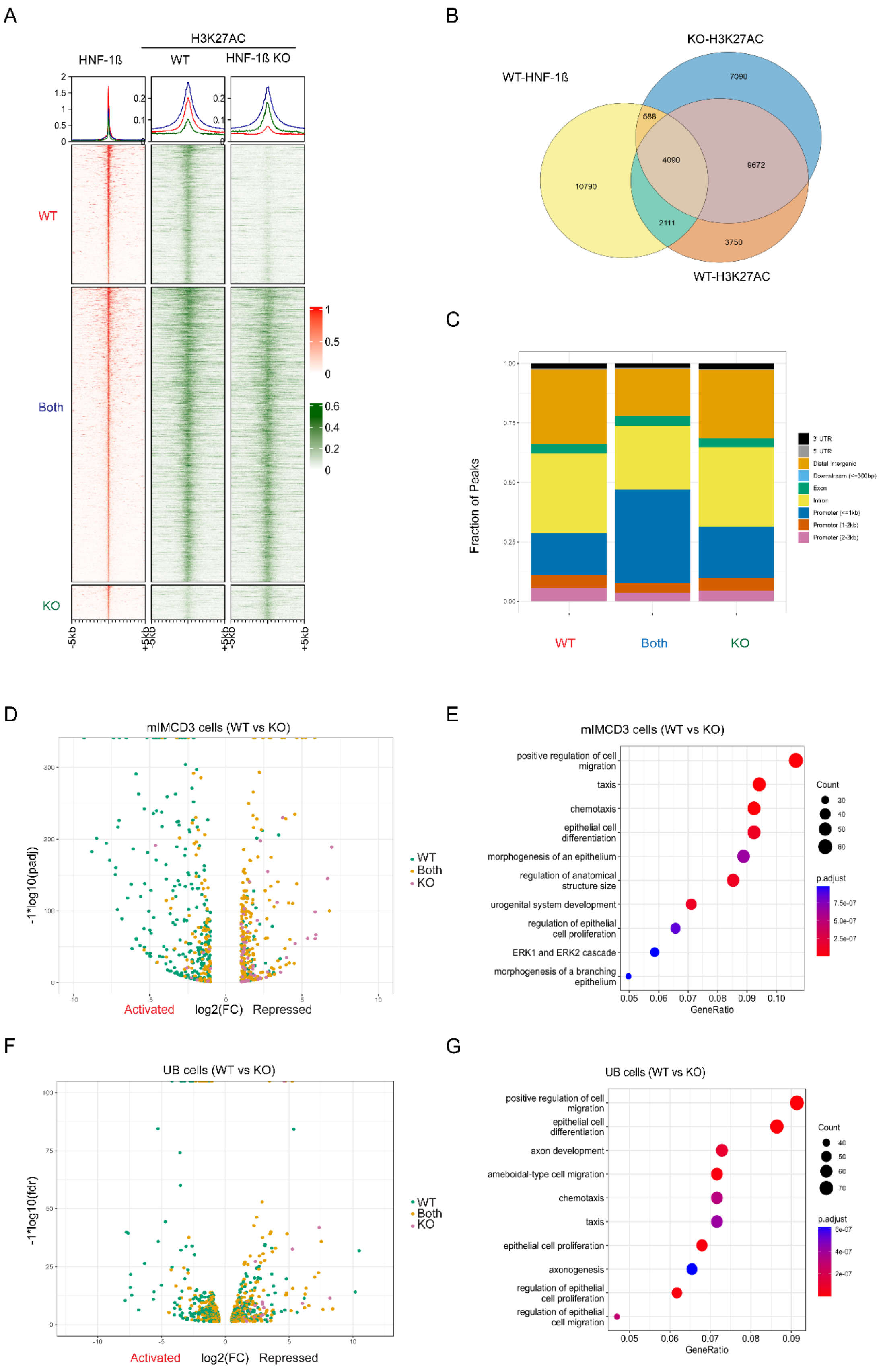

A
H3K27AC
HNF-1ß
WT
HNF-1ß KO
WT
Both
KO
-5kb
+5kb
B
KO-H3K27AC
WT-HNF-1ß
WT-H3K27AC
7090
588
4090
9672
10790
2111
3750
C
Fraction of Peaks
WT
Both
KO
D
mIMCD3 cells (WT vs KO)
-1*log10(padj)
Activated
log2(FC)
Repressed
WT
Both
KO
E
mIMCD3 cells (WT vs KO)
positive regulation of cell migration
taxis
chemotaxis
epithelial cell differentiation
morphogenesis of an epithelium
regulation of anatomical structure size
urogenital system development
regulation of epithelial cell proliferation
ERK1 and ERK2 cascade
morphogenesis of a branching epithelium
GeneRatio
Count
p.adjust
F
UB cells (WT vs KO)
-1*log10(fdr)
Activated
log2(FC)
Repressed
WT
Both
KO
G
UB cells (WT vs KO)
positive regulation of cell migration
epithelial cell differentiation
axon development
ameboidal-type cell migration
chemotaxis
taxis
epithelial cell proliferation
axonogenesis
regulation of epithelial cell proliferation
regulation of epithelial cell migration
GeneRatio
Count
p.adjust

**Figure S5. HNF-1β regulates histone H3K27 acetylation at distal enhancers. (**A) Heatmaps show ChIP-seq of HNF-1β (left lane) colocalized with histone H3K27 acetylation (H3K27ac) in wild-type mIMCD3 cells (WT) and HNF-1β-deficient cells (WT). Panel WT: Unique H3K27ac peaks colocalized with HNF-1β binding sites only in WT cells. Panel Both: Colocalized H3K27ac peaks at HNF-1β binding sites in the presence and absence of HNF-1β. Panel KO: Unique H3K27ac peaks only in KO cells. (B) Venn diagram shows the overlap between HNF-1β binding sites in WT cells and H3K27ac peaks in WT and KO cells. (C) Distribution of genomic locations of unique and shared H3K27ac peaks in the above three clusters (WT, Both and KO). (D) Volcano plots of genes that contained nearby colocalized HNF-1β and H3K27ac peaks and were differentially expressed in WT cells and KO cells. Genes with a nearby colocalized HNF-1β and H3K27ac peak only in wild-type mIMCD3 cells (green), in both wild-type and HNF-1β-deficient cells (gold), and only in HNF-1β-deficient cells (pink) are shown. (E) GO pathway analysis of HNF-1β-regulated genes located near colocalized HNF-1β and H3K27ac peaks. Dot size indicates the number of genes, and color indicates the adjusted p-value. (F) Volcano plots of genes that contained nearby colocalized HNF-1β and H3K27ac peaks and were differentially expressed in ureteric bud (UB) cells from E14.5 wild-type mouse kidneys (WT) and HNF-1β mutant kidneys (KO). (G) GO pathway analysis of differentially expressed genes located near colocalized HNF-1β and H3K27ac peaks in UB cells.

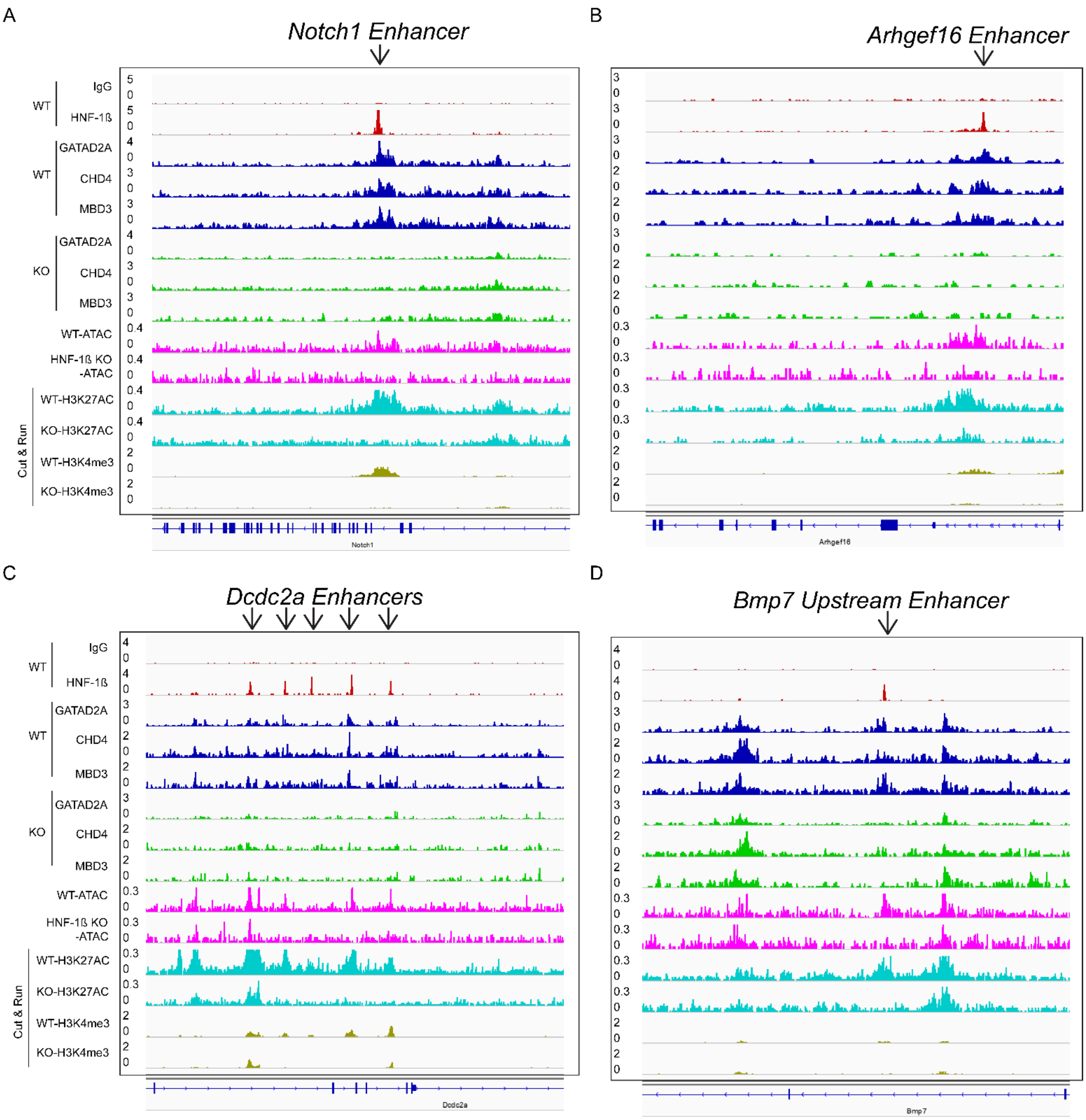


**Figure S6. HNF-1β recruits the NuRD complex resulting in opening of chromatin and upregulation of histone marks at active enhancers.** IGV gene tracks at Notch1 (A), Arhgef16 (B), Dcdc2a (C), and Bmp7 (D) showing ChIP-seq of HNF-1β, GATAD2A, CHD4 and MBD3; ATAC-seq; and CUT&RUN sequencing of H3K27ac and H3K4me3 in wild-type (WT) and HNF-1β-deficient (KO) mIMCD3 cells.

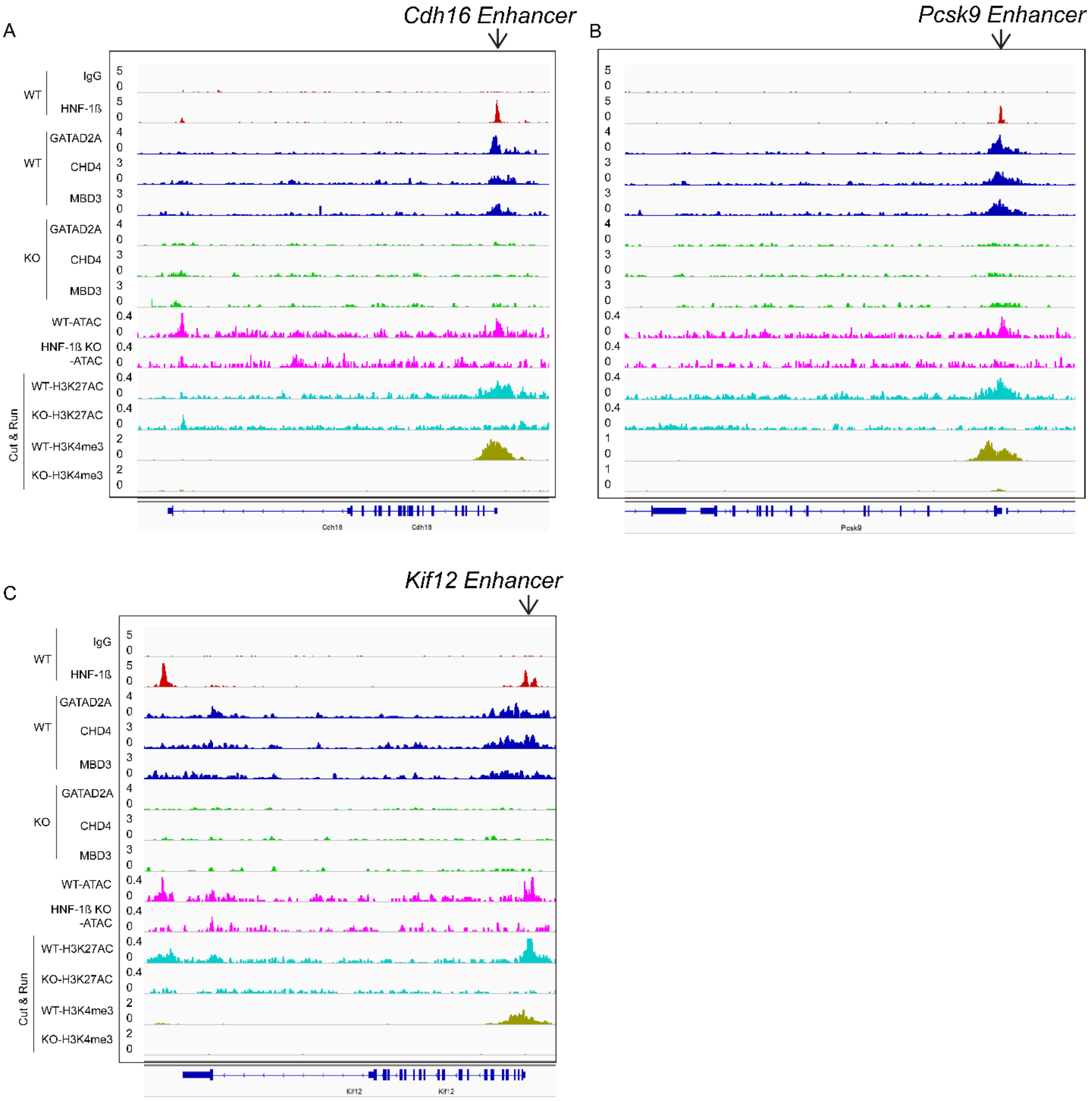


**Figure S7. HNF-1β-dependent recruitment of NuRD at enhancers of canonical HNF-1β target genes.** IGV gene tracks at canonical HNF-1b target genes Cdh16 (A), Kif12 (B), and Pcsk9 (C) showing ChIP-seq of HNF-1β, GATAD2A, CHD4 and MBD3; ATAC-seq; and CUT&RUN sequencing of H3K27ac and H3K4me3 in wild-type (WT) and HNF-1β-deficient (KO) mIMCD3 cells. Arrows indicate the positions of intronic enhancers exhibiting HNF-1β-dependent recruitment of NuRD, chromatin opening, H3K27 acetylation, and H3K4 trimethylation.
